# Highly accurate long-read HiFi sequencing data for five complex genomes

**DOI:** 10.1101/2020.05.04.077180

**Authors:** Ting Hon, Kristin Mars, Greg Young, Yu-Chih Tsai, Joseph W. Karalius, Jane M. Landolin, Nicholas Maurer, David Kudrna, Michael A. Hardigan, Cynthia C. Steiner, Steven J. Knapp, Doreen Ware, Beth Shapiro, Paul Peluso, David R. Rank

**Affiliations:** Pacific Biosciences of California Inc., 1305 O’Brien Dr., Menlo Park, CA 94025, USA; Ravel Biotechnology Inc., 953 Indiana St., San Francisco, CA 94107, USA; Department of Ecology and Evolutionary Biology, University of California Santa Cruz, Santa Cruz, CA 95064, USA; Arizona Genomics Institute and School of Plant Sciences, University of Arizona, Tucson, AZ 85721, USA; Department of Plant Sciences, University of California, Davis, One Shields Ave, Davis, CA 95616-8571, USA; Conservation Genetics, Beckman Center for Conservation Research, San Diego Zoo Global, 15600 San Pasqual Valley Road, Escondido, CA 92027, USA; Cold Spring Harbor Laboratory, Cold Spring Harbor, NY 11724, USA; USDA-ARS, Plant, Soil, and Nutrition Research Unit, Ithaca, NY 14853, USA; Howard Hughes Medical Institute, University of California Santa Cruz, Santa Cruz, CA 95064, USA

**Author notes:** Address correspondence to D.R.R.

## Abstract

The PacBio^®^ HiFi sequencing method yields highly accurate long-read sequencing datasets with read lengths averaging 10-25 kb and accuracies greater than 99.5%. These accurate long reads can be used to improve results for complex applications such as single nucleotide and structural variant detection, genome assembly, assembly of difficult polyploid or highly repetitive genomes, and assembly of metagenomes. Currently, there is a need for sample data sets to both evaluate the benefits of these long accurate reads as well as for development of bioinformatic tools including genome assemblers, variant callers, and haplotyping algorithms. We present deep coverage HiFi datasets for five complex samples including the two inbred model genomes *Mus musculus* and *Zea mays*, as well as two complex genomes, octoploid *Fragaria* × *ananassa* and the diploid anuran *Rana muscosa*. Additionally, we release sequence data from a mock metagenome community. The datasets reported here can be used without restriction to develop new algorithms and explore complex genome structure and evolution. Data were generated on the PacBio Sequel II System.

## Background & Summary

Until recently, DNA sequencing technologies produced either short highly accurate reads (up to 300 bases at 99% accuracy)^1,2^ or less-accurate long reads (10-100s of kb at 75-90% accuracy)^3,4^. Highly accurate short reads are appropriate for germline^5^ and somatic^6^ variant detection, exome sequencing^7^, liquid biopsy^8^, non-invasive prenatal testing^9^, and counting applications such as transcript profiling^10^ or single-cell analysis^11^. In contrast, error-prone long reads are more appropriate for *de novo* genome assembly^12–14^, haplotype phasing^15^, structural variant identification^16–18^, full-length mRNA sequencing and mRNA isoform discovery^19^.

To increase the utility of noisy long-read sequencing, several error correction methods have been devised to improve the accuracy of long reads by combining the data from either multiple independent long-read molecules or combining data from long-and short-read technologies^12,14^. These error-corrected reads can then be used for assembly or other downstream applications. In general, these error correction methods suffer from mis-mapping induced errors inherent to the multi-molecule approach^20^ that hinder downstream applications.

A third sequencing data type leveraging multiple pass circular consensus sequencing of long (>10 kb) individual molecules produces highly accurate long sequencing reads (HiFi reads)^21^. The HiFi sequencing protocol, data generation, and applications are described in **Figure 1**. In the initial publication^21^, 28-fold coverage of a human genome was sequenced with average read length of 13.5 kb and an average accuracy of 99.8%. The data has demonstrated superior assembly and haplotyping results for the human genome as measured by contiguity and accuracy when compared to traditional noisy long- or short-read methods. Additionally, single nucleotide variants were called at comparable precision and recall to Illumina^®^ NovaSeq^™^ data. Since the initial publication, greatly improved assembly results have been observed in other human sequencing projects^22–25^.

**Figure 1.**
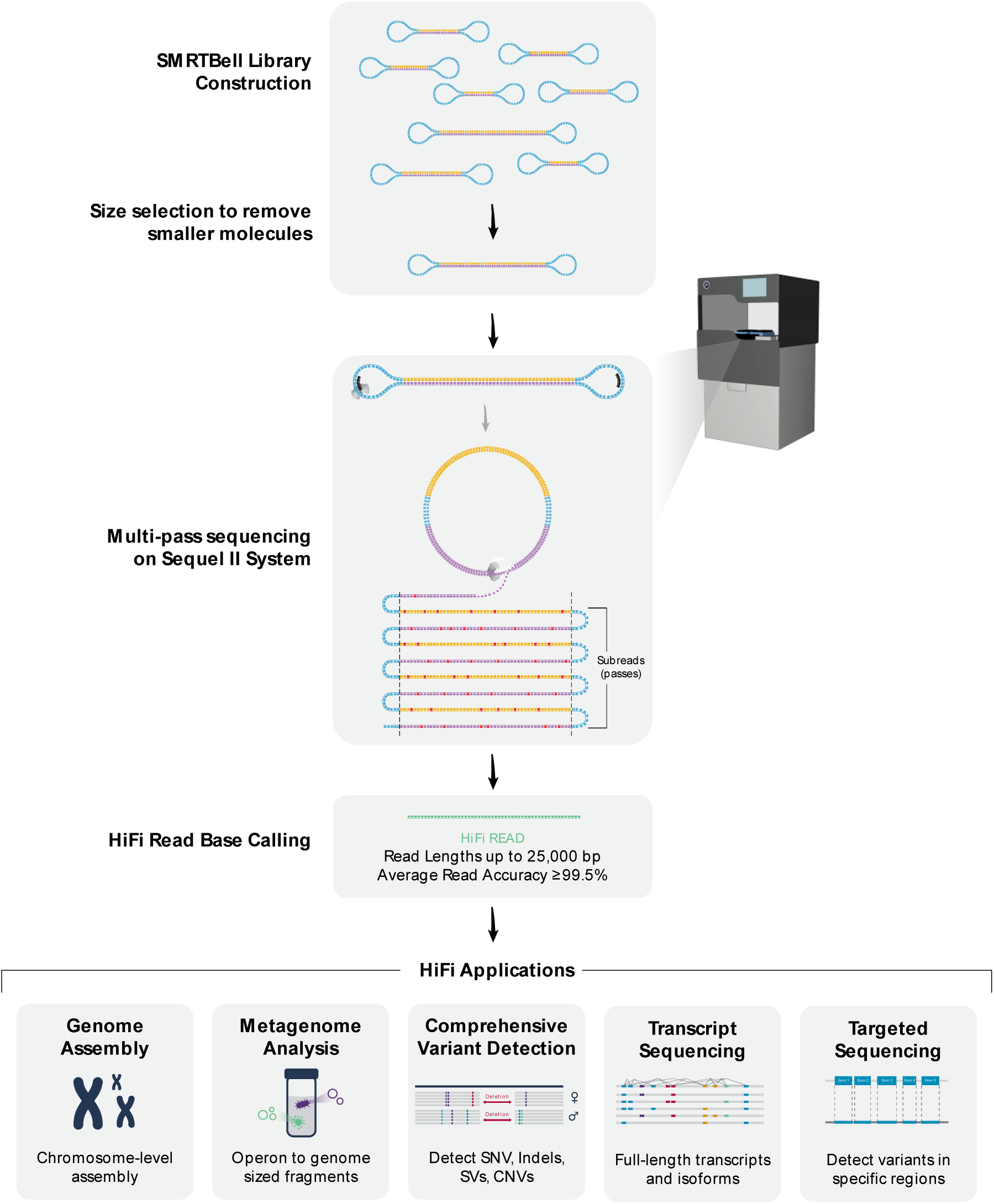
Flowchart of HiFi sequence read generation and downstream applications.

To encourage further application, software development, and interest in the HiFi sequencing data type, we report here the release of five deep coverage data sets spanning a set of complex genomes including *Mus musculus, Zea mays, Fragaria* × *ananassa* (Weston) Duchesne ex Rozier, *Rana muscosa*, and a standard metagenomic collection of 20 microbes formulated at staggered concentrations (ATCC^®^ MSA-1003^™^). The data released in this study covers a wide breadth of highly complex plant, animal, and microbial organisms and will provide a useful sequence resource, driving the sequencing standards toward higher quality in the future^25^.

## Methods

### Sample Selection

Organisms sequenced in this study include *M. musculus, Z. mays, F.* × *ananassa*, and *R. muscosa*. The strain of each organism, source of the material, ploidy level, inbreeding status, reference genome sequence, and genome sizes are described in **Tables 1 and 2**. Additionally, we are releasing sequencing reads from a mock metagenomic sample (ATCC MSA-1003) consisting of 20 bacterial DNA samples at staggered concentrations ranging from 0.02% to 18% composition of the sample. The composition of the mock metagenomic sample as well as genome sizes of the individual bacterial species and their reference sequence accessions are listed in **Supplementary Table 1**.

**Table 1.** Sample description: strain names, origins, available reference sequences, and SRA Bio Sample IDs are detailed for each HiFi dataset.

| Sample | Strain<br>(Cultivar/ Cell<br>line) | Sample<br>Origin | Sequence Reference | SRA Bio<br>Sample ID |
| --- | --- | --- | --- | --- |
| ***M. musculus*** | C57BL/6J | Jackson<br>Labs | GRCm38.p6* | SAMN14691541<sup>#</sup> |
| ***Z. mays*** | B73 | M. Hufford | Zm-B73-REFERENCE-<br>NAM-5.0<sup>‡</sup> | ERS3371164<sup>§</sup><br>SAMN14691542<sup>#</sup> |
| ***F. x ananassa*** | Royal Royce | S. Knapp | N/A | SAMN14691544<sup>#</sup> |
| ***R. muscosa*** | KB 21384; ISIS<br># 916035 | San Diego<br>Zoo Global | N/A | SAMN14691543<sup>#</sup> |
| **Metagenome Std** | MSA-1003 | ATCC | See **Supplementary**<br>**Table 1** | SAMN14691545<sup>#</sup> |

\*https://www.ncbi.nlm.nih.gov/assembly/GCF_000001635.26/
<sup>#</sup>https://www.ncbi.nlm.nih.gov/bioproject/PRJNA627939
<sup>‡</sup>https://www.ncbi.nlm.nih.gov/assembly/GCA_902167145.1/
<sup>§</sup>https://www.ncbi.nlm.nih.gov/bioproject/600450

**Table 2.**
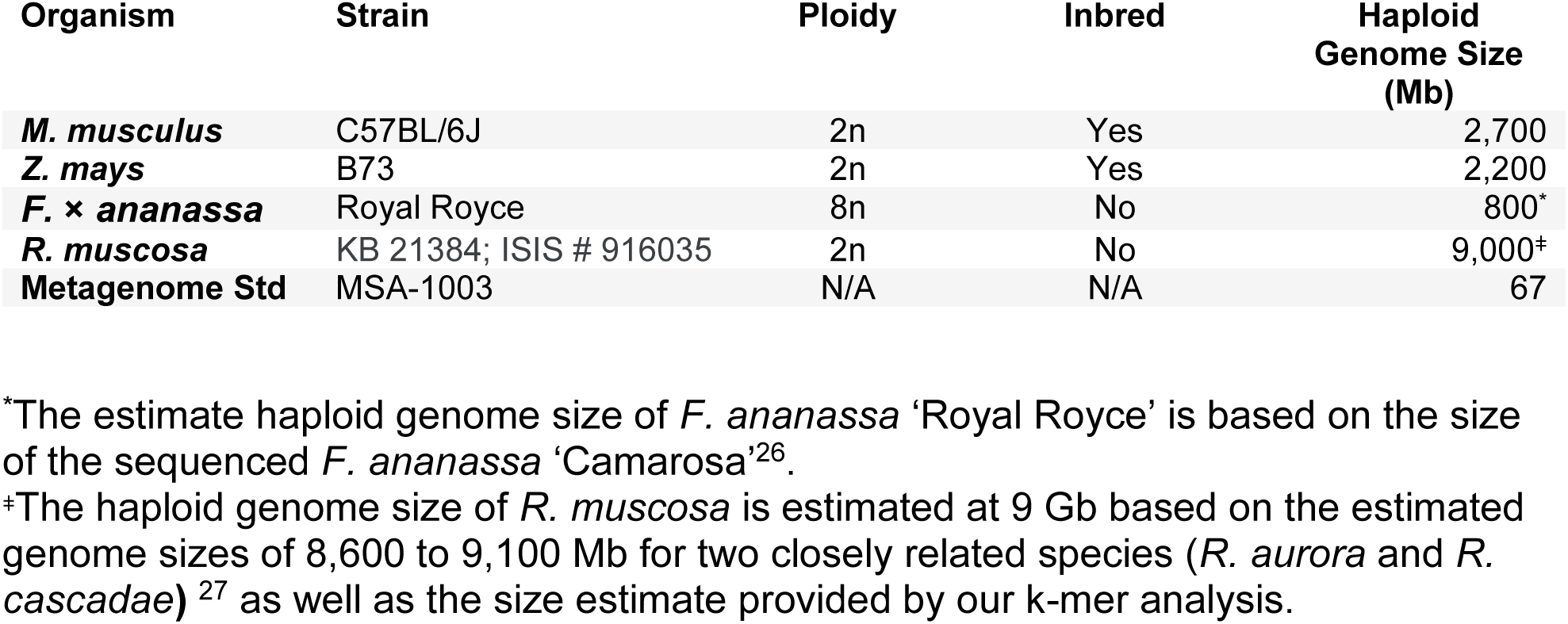
Background genomic information for each sample: strain or sample ID, expected ploidy level, inbred status, and haploid genome size for each HiFi read dataset.

Excluding the metagenomic sample, the expected assembly sizes for the genomes sequenced in this study ranged from the 1,600 Mb for the outbred and octoploid *F.* × *ananassa*^26^ to approximately 18,000 Mb for the outbred and diploid *R. muscosa (*estimate based on genome sizes of two related species *Rana aurora* and *Rana cascadae)*^27^. The individual genome sizes of the metagenomic sample range from 1.67 to 6.34 Mb, totaling 67 Mb of bacterial sequence (**Supplementary Table 1**).

### Sequencing Library Preparation

Genomic DNA extraction methods and details of individual library preparations are described in the sample specific sections below. In general, if the starting genomic DNA sample was larger than 25 kb, the DNA was sheared to between 15 kb and 23 kb using the Megaruptor^®^ 3 (Diagenode). HiFi sequencing libraries were prepared^28^ using SMRTbell™ Express Template Prep Kit 2.0 and followed by immediate treatment with the Enzyme Clean Up Kit (PN: 101-843-100). The libraries were further size selected electrophoretically using either the SageELF or BluePippin Systems from SAGE Science. The appropriate fractions for sequencing runs were identified on the Femto Pulse System (Agilent). After pooling the desired size fractions, the final libraries were further cleaned up and concentrated using AMPure PB beads (Pacific Biosciences PN:100-265-900). Finally, all libraries were checked for concentration using Qubit™ 1X dsDNA HS Assay Kit (Thermo Fisher PN: Q33231) and final size distribution was confirmed on the Femto Pulse. All library sizes are described in **Table 3**.

**Table 3.**
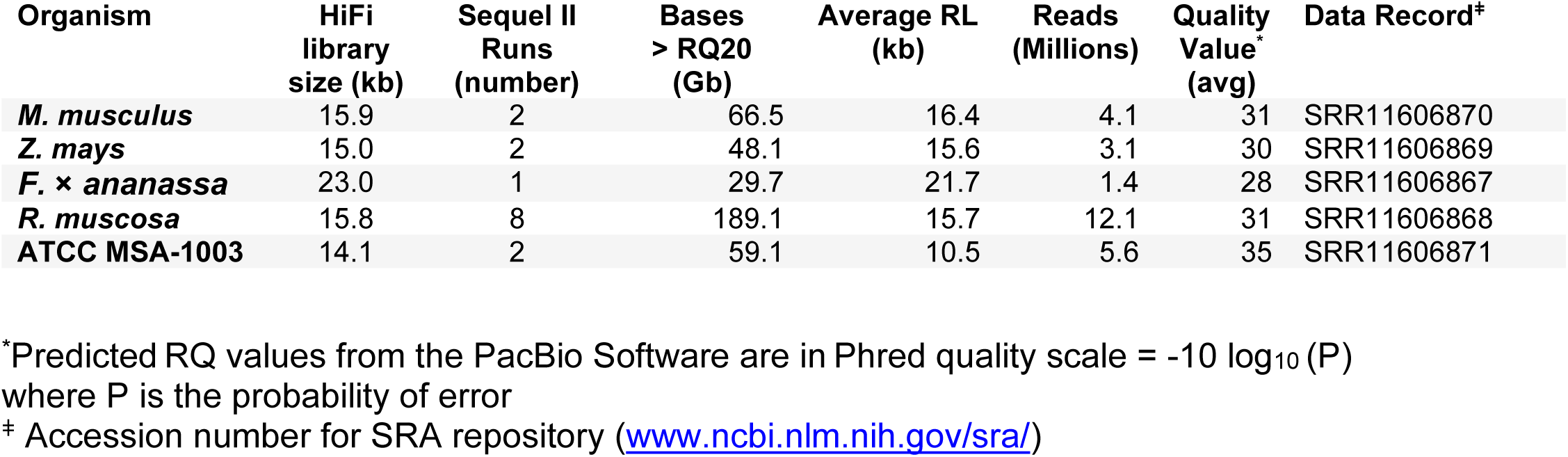
Library molecule sizes, sequencing metrics, and SRA accession numbers for each HiFi read dataset.

| Organism | HiFi library size (kb) | Sequel II Runs (number) | Bases > RQ20 (Gb) | Average RL (kb) | Reads (Millions) | Quality Value* (avg) | Data Record<sup>‡</sup> |
| --- | --- | --- | --- | --- | --- | --- | --- |
| *M. musculus* | 15.9 | 2 | 66.5 | 16.4 | 4.1 | 31 | SRR11606870 |
| *Z. mays* | 15.0 | 2 | 48.1 | 15.6 | 3.1 | 30 | SRR11606869 |
| *F. x ananassa* | 23.0 | 1 | 29.7 | 21.7 | 1.4 | 28 | SRR11606867 |
| *R. muscosa* | 15.8 | 8 | 189.1 | 15.7 | 12.1 | 31 | SRR11606868 |
| ATCC MSA-1003 | 14.1 | 2 | 59.1 | 10.5 | 5.6 | 35 | SRR11606871 |

\*Predicted RQ values from the PacBio Software are in Phred quality scale = $-10 \log_{10}(P)$ where P is the probability of error
<sup>‡</sup> Accession number for SRA repository (www.ncbi.nlm.nih.gov/sra/)

### *M. musculus* ‘C57/BL6J’ sample acquisition, DNA extraction, and modifications to sequencing library preparation

C57/BL6J genomic DNA was obtained from The Jackson Laboratory (PN: GTC4560). The DNA arrived at an appropriate size for HiFi library preparation (∼20 kb) and no shearing was required. Library preparation method, kit, and conditions were as described above. In order to tighten the size distribution of the SMRTbell library, the DNA was size fractionated using the SageELF following library preparation. The SMRTbell Library was prepared with loading solution/Marker75 then loaded onto a 0.75% agarose 1kb-18 kb gel cassette (PN: ELD7510). Size fractionation was performed electrophoretically with a target size of 3,500 bp set for elution well 12, which allowed for the collection of the appropriately sized library fractions (15-23 kb) in other elution wells of the SageELF device.

### *Z. mays* ‘B73’ sample acquisition, DNA extraction, and modifications to sequencing library preparation

Leaf tissue for the B73 maize inbred was frozen and provided by Matthew Hufford at Iowa State University, Department of Ecology, Evolution, and Organismal Biology. Genomic DNA was isolated from the frozen leaf tissue at the University of Arizona Genomics Institute using methods previously described^29^. The high molecular weight DNA was sheared using the Megaruptor 3 targeting a size distribution between 15 and 20 kb. Library preparation method, kit and conditions were as described above. Library size selection was performed on the Sage BluePippin using the 0.75% Agarose dye-free Gel Cassette (PN: BLF7510) and the S1 Marker. To ensure suitable yields, the 3-10 kb Improved Recovery cassette definition was run for the size selection and high pass elution mode was chosen to target recovery of molecules greater than 15 kb.

### *F.* × *ananassa* ‘Royal Royce’ sample acquisition, DNA extraction, and modifications to sequencing library preparation

The plant material was obtained from foundation stock of the cultivar ‘Royal Royce’ maintained by the UC Davis Strawberry Breeding Program. DNA was isolated as previously described^30^. The genomic DNA was larger than required for HiFi library production and was sheared using the Megaruptor 3 targeting a size distribution centered around 22 kb. Library preparation method, kit, and conditions were as described above. The SageELF was used for size selection, with similar conditions as described for *M. musculus* above, in order to generate a library with an appropriately sized distribution.

### *R. muscosa* sample acquisition, DNA extraction, and modifications to sequencing library preparation

*R. muscosa*, the Mountain Yellow-legged Frog, is an endangered species endemic to California. To prevent sacrificing an individual, DNA was prepared from a fibroblast cell line (KB 21384; ISIS # 916035) originally derived from a 25-day old tadpole of undetermined sex. The cells were grown at room temperature in low O_2_ from explants in alpha MEM with 1% NEAA. Approximately two million cells were harvested at passage 7 and frozen in a 1X solution of PBS buffer with 10% DMSO and 10% glycerol.

Genomic DNA was isolated from these cells using Qiagen’s MagAttract HMW DNA Kit (PN: 67563) following the manufacture’s protocol. The resulting HMW gDNA was sheared to a target size of 22 kb on the MegaRuptor 3 prior to library preparation. Library preparation, kit and conditions were as described above. In order to tighten the size distribution, the SMRTbell library was size fractionated using SageELF System from Sage Science. The DNA was premixed with loading solution/Marker40 and loaded onto a 0.75% Agarose 10-40 kb Cassette (PN: ELD4010). Size fractionation was performed electrophoretically with a target size of 7,000 bp set for elution well 12 in order to achieve the appropriate resolution in size separation. Fractions having the desired size distribution ranges were identified on the Femto Pulse to generate a final size selected library used in the Sequel II sequencing runs. An additional DNA damage repair step was performed using the SMRTbell Damage Repair Kit (PN:100-992-200) as this was found helpful to improve library performance in sequencing runs.

### Mock metagenome sample acquisition, DNA extraction, and modifications to sequencing library preparation

ATCC offers a mock metagenomic community (MSA 1003) of 20 bacteria species ranging in composition from 0.02% to 18% of the sample. Isolated DNA from this sample arrived with genomic DNA having a broad distribution of sizes and was sheared using the MegaRuptor 3 to a uniform size of 13.7 kb. Library preparation method, kit and condition were described above. Rather than using electrophoretic size selection, the resulting library was size selected using AMPure PB beads (35% v/v) to remove all small fragments.

### Sequencing and Data Processing

SMRTbell libraries were bound to the sequencing polymerase enzyme using the Sequel II Binding Kit 2.0 (PN:101-842-900) with the modification that the Sequencing Primer v2 (PN:101-847-900) was annealed to the template instead of the standard primer which comes with Sequel II Binding Kit 2.0. All incubations were performed per manufacturer’s recommendations. Prior to sequencing, unbound polymerase enzyme was removed using a modified AMPure PB bead method as previously described^21,31^. Shotgun genomic DNA sequence data was collected on the Pacific Biosciences Sequel II system using HiFi sequencing protocols^31^ and Sequencing kit V2 (PN: 101-820-200). Sequence data collection was standardized to 30 hours for this study to allow ample time for multiple pass sequencing around SMRTbell template molecules of 10-25 kb which yields high quality circular consensus sequencing (HiFi) results^21^. Raw base-called data was moved from the sequencing instrument and the imported into SMRTLink^32^ to generate HiFi reads using the CCS algorithm (version 8.0.0.80529) which processed the raw data and generated the HiFi fastq files with the following settings: minimum pass 3, minimum predicted RQ 20.

### K-mer Analysis

Using Jellyfish^33^ (v.2.2.10) a k-mer analysis was performed on each of the HiFi data sets individually using a k-mer size of 21. Counting was done using a two-pass method. First, a Bloom counter was created for each HiFi read dataset using the following command:

~~~
jellyfish bc -m 21 -s <Input Size> -t <nproc> -C -o
HiFiReadSetFilename.bc HiFiReadSet.fasta
~~~

where Input Size = 100G (*M. musculus, Z. mays, F.* × *ananassa and R. muscosa*) and 5G (ATCC MSA-1003).

After generating the Bloom counter, a frequency count of k-mers (size=21) was run using the following command:

~~~
jellyfish count -m 21 -s <Input Size> -t <nproc> -C --bc
HiFiReadSetFilename.bc HiFiReadSet.fasta
~~~

Where Input Size = 20G (*R. muscosa*), 3G (*M. musculus* and *Z. mays*), 2G (*F.* × *ananassa*) and 200M (ATCC MSA-1003).

Finally, a histogram of the k-mer frequency was generated for each dataset using the following command:

~~~
jellyfish histo HiFiReadSet_21mer counts.jf >
HiFiReadSet_21mer_Histogram.out
~~~

These outputs were then used to generate the additional summary analysis and determine genome sizes for each sample where applicable. Genome sizes were estimated from the ratio of total HiFi bases divided by the frequency mode from each k-mer distribution.

### Mapping Accuracies and Read Lengths

In the cases where references were available (*M. musculus, Z. mays*, and the concatenated genomes comprising the ATCC MSA-1003 sample), HiFi reads were mapped to the references using pbmm2 version 1.2.0 (https://github.com/PacificBiosciences/pbmm2) which is a customized wrapper for minimap2^34^ using the following command:

~~~
pbmm2 align REF.fasta HiFiReadSet.fastq
HiFiReadSet.REF.sorted.bam --preset CCS --sort -j 48 -J 16
~~~

(where j + J= nproc=64)

To extract accuracy metrics from each bam file using Samtools^35^ version 1.9, the following command was used:

~~~
samtools view HiFiReadSet.REF.sorted.bam | awk ‘{ mc=““;
for(i=12;i<=NF;i++) { split($i,TAG,”:”); if(TAG[1]==“mc”) {
mc=TAG[3]; break; } } if(mc != ““) { print $1 “\t” mc; } }’ >
MappedConcordance.HiFiReadSet.Genome.out
~~~

To extract read length metrics from each bam file using Samtools, the following command was used:

~~~
samtools view HiFiReadSet.REF.sorted.bam | head -n <input # of
HiFi Reads> | cut -f 10 | perl -ne ‘chomp;print length($_).
“\n”’ | sort | uniq -c > MappedRL.HiFiReadSet.Genome.out
~~~

Finally, coverage metrics were obtained from each bam files using the following command in Samtools:

~~~
samtools depth -a HiFiReadSet.REF.sorted.bam >
HiFiReadSet.REF.sorted.Depth.out
~~~

### Data Records

All sequencing data presented are available at the Sequencing Read Archive (SRA) under BioProject accession number PRJNA627939. The HiFi sequencing data is stored as fastq files with one file for each Sequel II sequencing run. Information describing each data record is presented in **Table 3** and described below.

**SRR11606870**^36^ The *M. musculus* ‘C57/BL6J’ data record is composed of two Sequel II runs (total of two SMRT Cell 8M) containing 4.1 M sequencing reads and 66.5 Gb of sequence which corresponds to 25-fold coverage of the mouse genome. The average read length is 16.4 kb with an average PacBio predicted quality value (RQ) of 31.

**SRR11606869**^37^ The *Z. mays* ‘B73’ data record is composed of two Sequel II runs (total of two SMRT Cell 8M) containing 3.1 M sequencing reads and 48.1 Gb of sequence which corresponds to 22-fold coverage of the maize genome. The average read length is 15.6 kb with an average PacBio predicted quality value of 30.

**SRR11606867**^38^ The *F.* × *ananassa* ‘Royal Royce’ data record is composed of one Sequel II run (total of one SMRT Cell 8M) containing 1.4 M sequencing reads and 29.7 Gb of sequence of the octoploid Royal Royce genome. The average read length is 21.7 kb with an average RQ value of 28.

**SRR11606868**^39^ The *R. muscosa* (cell line KB 21384; ISIS # 916035) data record is composed of 8 Sequel II runs (total of eight SMRT Cell 8M) containing 12.1 M sequencing reads and 189.1 Gb bases of sequence which corresponds to approximately 20-fold coverage of the *R. mucosa* genome. The average read length is 15.7 kb with an average RQ of 31.

**SRR11606871**^40^ is the data record for the ATTC MSA-1003 mock metagenome community which is composed of 20 bacterial organisms reported to be mixed at relative amounts differing by 900 fold from highest to lowest (**Supplementary Table 1**). The files in this sequence record span two Sequel II runs (total of two SMRT Cell 8M) containing 5.6 M sequencing reads with 59.1 Gb of sequence which corresponds to between ∼3 and ∼5,000-fold coverage of the individual bacterial genomes. The average read length is 10.5 kb with an average RQ value of 35.

### Technical Validation

Two of the non-microbial organisms sequenced, *M. musculus*, and *Z. mays*, have high quality reference genomes available^41,42^ allowing for detailed validation of the sequencing data. Additionally, reads from sequencing the mock metagenome sample were aligned to a concatenated file containing all microbial references listed in **Supplementary Table 1** and used for validation. All reads were aligned to their corresponding references using pbmm2. The read accuracy and mapped read length are reported in **Table 4** and distributions are presented in **Figure 2**. In agreement with previously published reports^21^, the accuracy of the HiFi reads exceeds 99.5% (median accuracy of 99.87%, 99.84%, and 99.99% for the mouse, maize and mock metagenome samples respectively). The mean accuracies were 99.18% (mouse), 99.69% (maize) and 99.73% (mock metagenome). Sequencing read lengths (**Table 3**) ranged from 10.5 kb (mock metagenome) to 21.7 kb (*F.* × *ananassa*) and were dependent on the final size distributions of the sequencing libraries.

**Table 4.** Technical validation summary: k-mer based genome size estimates, average mapped HiFi read coverage, and average mapped HiFi read accuracy for each dataset.

| Sample | K-mer based<br>Genome<br>Coverage (fold) | Reference<br>Mapped Genome<br>Coverage (fold) | Median Read<br>Accuracy<br>(percent) | Mean Read<br>Accuracy<br>(percent) |
| --- | --- | --- | --- | --- |
| *M. musculus* | 25 | 27<sup>*</sup> | 99.869 | 99.176 |
| *Z. mays* | 21 | 23<sup>‡</sup> | 99.844 | 99.686 |
| *F. x ananassa* | 17/37/74/109 | N/A<sup>#</sup> | N/A<sup>#</sup> | N/A<sup>#</sup> |
| *R. muscosa* | 20 | N/A<sup>#</sup> | N/A<sup>#</sup> | N/A<sup>#</sup> |
| ATCC MSA-1003 | 2 - 4000 | 1 - 8,000<sup>§</sup> | 99.995 | 99.733 |

<sup>\*</sup>https://www.ncbi.nlm.nih.gov/assembly/GCF_000001635.26/
<sup>‡</sup>https://www.ncbi.nlm.nih.gov/assembly/GCA_902167145.1/
<sup>#</sup> No published reference yet available
<sup>§</sup> See **Supplementary Table 1** for reference genome file names and locations

**Figure 2.**
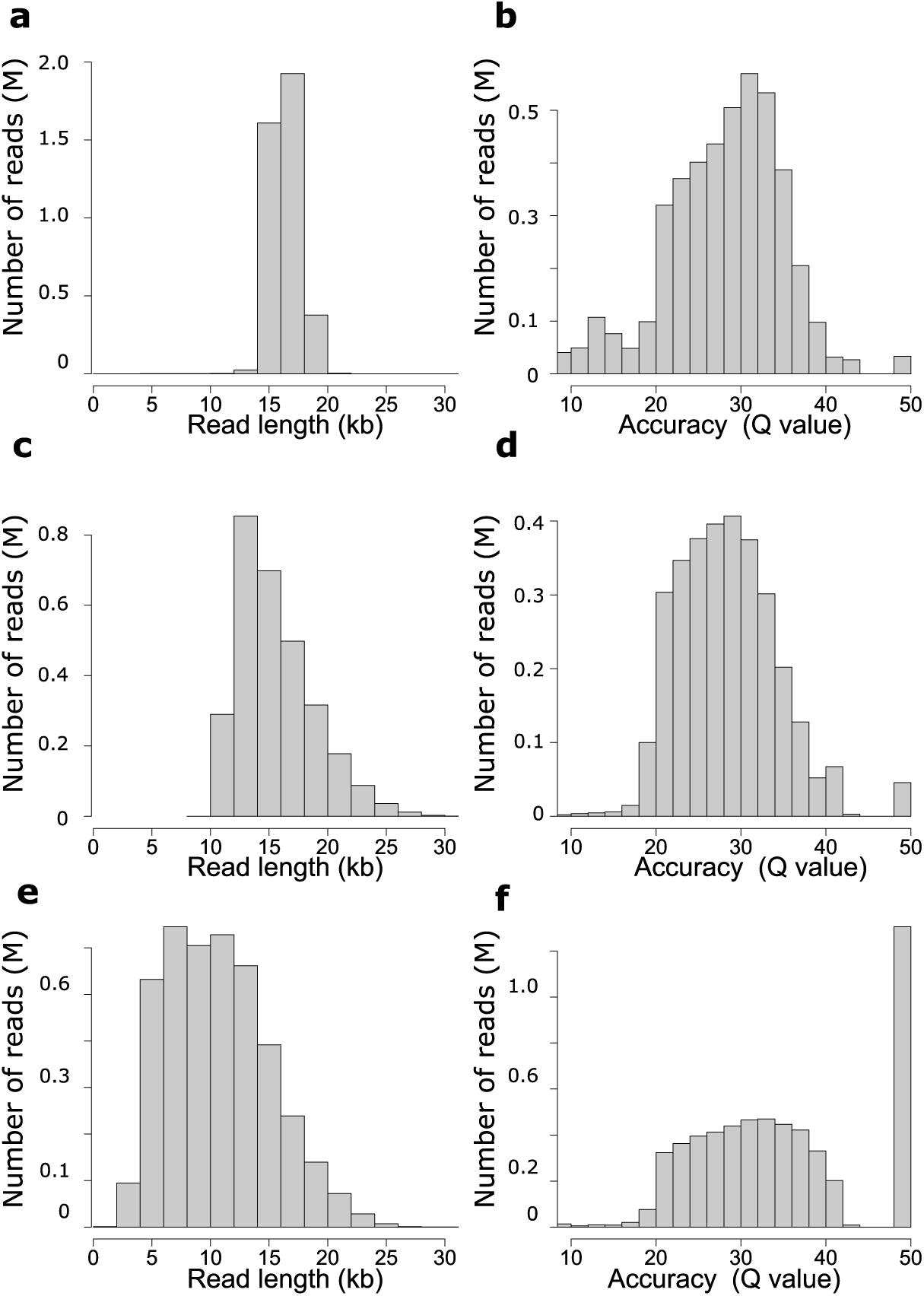
Read length and quality distributions for the three sequenced samples with high quality finished sequence references. *M. musculus* read length (a) and accuracy (b), *Z. mays* read length (c) and accuracy (d), and Mock metagenome community ATTC MSA-1003 read length (e) and accuracy (f). All data is mapped to the genomic references (**Table 1 and Supplementary Table 1)** using minmap2. Accuracies are reported in Phred read quality space (Q value) = −10 × log_10_(P) where P is the measured error rate.

The data for all five organisms was used to generate k-mer plots using a k-mer size of 21 (**Figure 3 a-e**) to estimate the sequencing coverage and complexity of for each sample. K-mer based sequencing coverage was measured at 17 to 25-fold (**Table 4**) for each of the individual diploid genomes sequenced (*M. musculus, Z. mays, and R. muscosa*) and as expected produced a multimodal distribution for the octoploid *F.* × *ananassa* and a complex curve for the metagenome sample.

**Figure 3.**
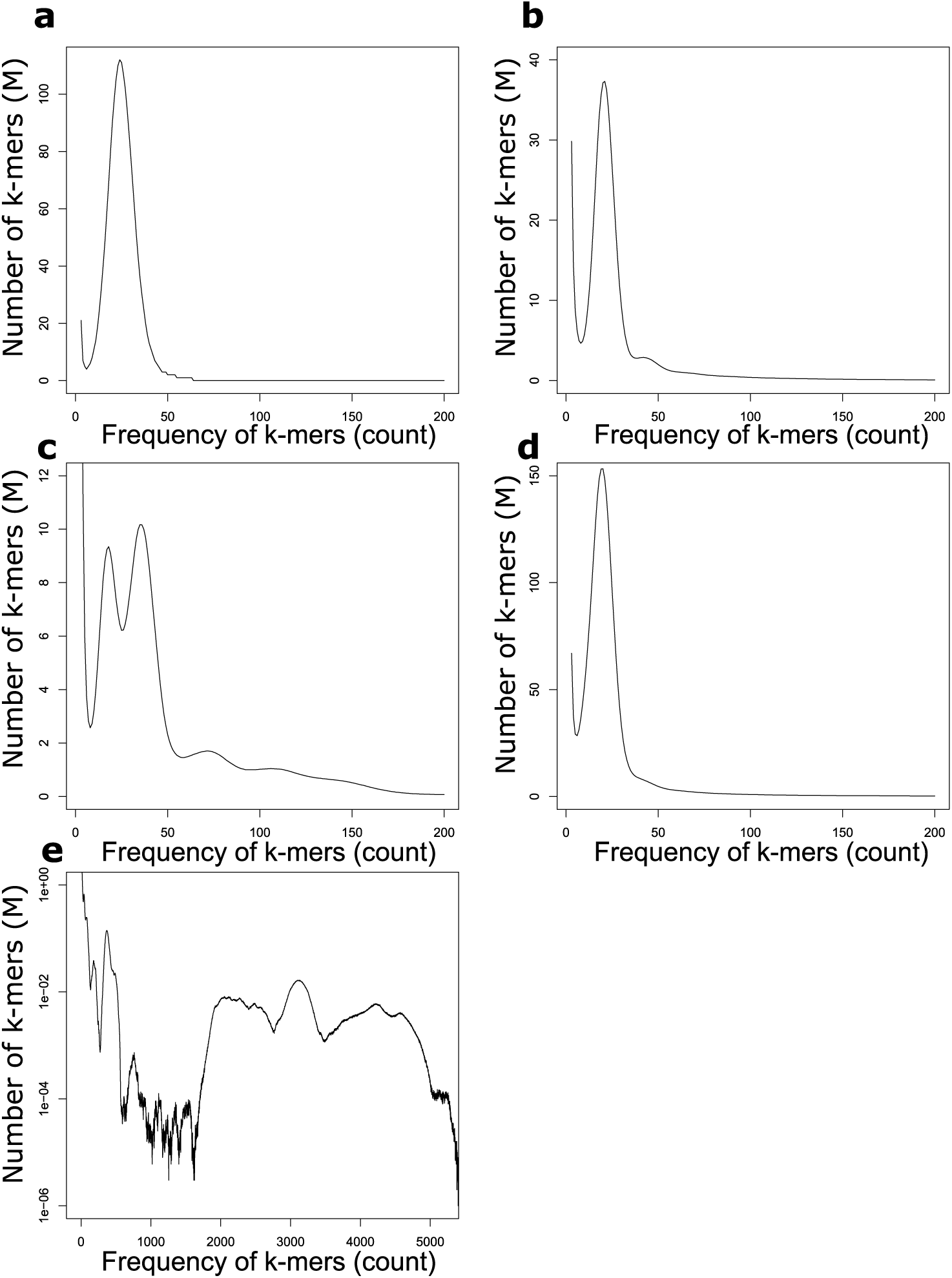
K-mer (length 21) distribution for all HiFi reads for each sequencing dataset. (a) *M.* musculus (b) *Z. mays* (c) *F.* × *ananassa* (d) *R. muscosa* (e) Mock metagenome community ATTC MSA-1003.

Additionally, the k-mer plots can be used to characterize genome complexity such as ploidy and/or genome duplications as evidenced by multimode distributions within the k-mer plots caused by inherent polymorphism within the respective genomes. As expected, the inbred mouse C57/B6J, shows a single k-mer distribution consistent with the single haplotype present in the inbred animal. The inbred B73 maize shows a dominant k-mer coverage peak at 21-fold as one would expect, but also a minor peak at 42-fold which is consistent with an ancient duplication and polyploidization^43^ of this inbred sample. The octoploid strawberry *F.* × *ananassa* has a k-mer plot consistent with the known complexity of this polyploid sample, having k-mer peaks, at 17, 37, 74, and 109.

The diploid *R. muscosa* sample demonstrates a more interesting case with respect to k-mer analysis as the frequency distribution shows one single haplotype at 20-fold coverage. The presence of a single k-mer peak in the genomic reads likely speaks to population bottlenecking which reduced the level of polymorphism in the genome resulting in collapse of the paternal and maternal haplotypes into one frequency peak for a k-mer size of 21. This is further supported by an apparent haploid genome size of 9,000 Mb (as calculated by the total number of sequenced bases / frequency mode of the k-mer histogram) which is equal to one half the size of the measured diploid genome sizes (∼18 Gb) of two closely related *species* (*R. aurora* and *R. cascadae*)^27^.

Alternatively, the genome coverage can be measured by mapping the HiFi reads to published references. The genome wide mapping-based coverages are reported in **Table 4** and agree with the k-mer based estimates for those samples with known references. Average mapped coverage values for each chromosome for both *M. musculus* and *Z. mays* are reported in **Supplementary Table 2** and demonstrate the even coverage distribution for the samples. The mapping method for genome coverage calculation also produces coverage values for each member of the mock metagenome community sample and is consistent with the genomic complexity displayed in the k-mer plot (**Figure 3e**), and agrees with the uneven representation of the abundance of each microbe in the mixture (**Supplementary Figure 1**).

### Usage Notes

The data presented in this manuscript should provide ample DNA sequence for genome assembly, variant detection, evaluation of metagenome completeness and metagenome assembly for the samples covered. Additionally, the data should prove useful for bioinformaticians developing, improving, and validating assembly algorithms, developing haplotyping tools, and variant detection algorithms. High contiguity and high-quality genome assemblies should also be possible for the two unpublished genomes presented in this study (*F.* × *ananassa* ‘Royal Royce’, and the endangered anuran *R. muscosa)*. Recently, HiFi read based assemblies have reconstructed several centromeres of the human genome^25^, and the HiFi data presented here will be useful for future updates of the reference genomes for both *Z. mays* ‘B73’ and *M. muscosa* ‘C57/BL6J’ by adding previously unresolvable regions, possibly including some complete centromeres of these genomes. The data from the metagenome mock community should prove valuable for metagenome assembly algorithms, and other analytical tool development allowing for the assembly of complete bacterial genomes from metagenomic samples displaying high heterogeneity in individual bacterial species and relative abundance.

## Code Availability

Bioinformatic tools used for validation are all open source and feely available. We used jellyfish version 2.2.10 to count k-mers (https://github.com/gmarcais/Jellyfish), pbmm2 version 1.2.0 to map to a reference (https://github.com/PacificBiosciences/pbmm2), and samtools version 1.9 to summarize metrics (http://www.htslib.org).

## Acknowledgements

The contributions of S.J.K. were funded by grants from the United Stated Department of Agriculture National Institute of Food and Agriculture (NIFA) Specialty Crops Research Initiative (2017-51181-26833), California Strawberry Commission, and the University of California. The contributions of D.W. were funded by USDA-ARS 8062-21000-041, NSF IOS-1744001.

We would like to thank Mathew Hufford and Kelly Dawe for providing the B73 maize leaf material as well as insightful discussions on the manuscript.

We would like to thank Marlys Houck and Catherine Avila for their important contributions in producing the cell line used to generate sequence of the Mountain Yellow-legged Frog *R. muscosa*.

Additionally, we thank Kristin Robertshaw and Pamela Bentley Mills for technical assistance generating figures.

## Author contributions

T.H., K.M., G.Y., and Y-C.T. library preparation, DNA sequencing and data quality control, and manuscript preparation. J.M.L. and J.W.K. collating sequencing data, posting to data repositories, and manuscript preparation. N.M., D.K., M.A.H. sample selection, sample preparation, and DNA isolation. C.C.S., S.J.K., D.W., and B.S. sample selection and manuscript preparation. P.S.P. experimental design, sequencing coordination, data submission, technical validation, bioinformatic analysis, and manuscript preparation. D.R.R. experimental design, technical evaluation, and manuscript preparation.

## Competing Interests

T.H., K.M., G.Y., Y-C. T., J.W.K., P.S.P., D.R.R. are employees of Pacific Biosciences of California Inc. a company commercializing DNA sequencing technology.

J.M.L. is an employee of Ravel Biotechnology Inc. a company commercializing disease detection from cell-free DNA.

All other authors declare no competing interests.

## Notes

https://identifiers.org/ncbi/insdc.sra:SRR11606870

https://identifiers.org/ncbi/insdc.sra:SRR11606869

https://identifiers.org/ncbi/insdc.sra:SRR11606867

https://identifiers.org/ncbi/insdc.sra:SRR11606868

https://identifiers.org/ncbi/insdc.sra:SRR11606871

