## Supplementary Material for "Highly accurate long-read HiFi sequencing data for five complex genomes"

### Detailed Author Contributions

Library preparation and DNA sequencing, data quality control, and manuscript preparation: T.H., K.M., G.Y., and Y-C.T.

Collated sequencing data and posted to data repositories and manuscript preparation: J.M.L, J.W.K.

Sample preparation and DNA isolations: N.M., D.K., M.A.H.

Sample selection and manuscript preparation: C.C.S., S.J.K., D.W., B.S.

Experimental design, sequencing coordination, data submission, performed technical validation, bioinformatic analysis, and manuscript preparation: P.S.P.

Experimental design, technical evaluation, and manuscript preparation: D.R.R.

### Supplementary Figures

Supplementary Figure 1

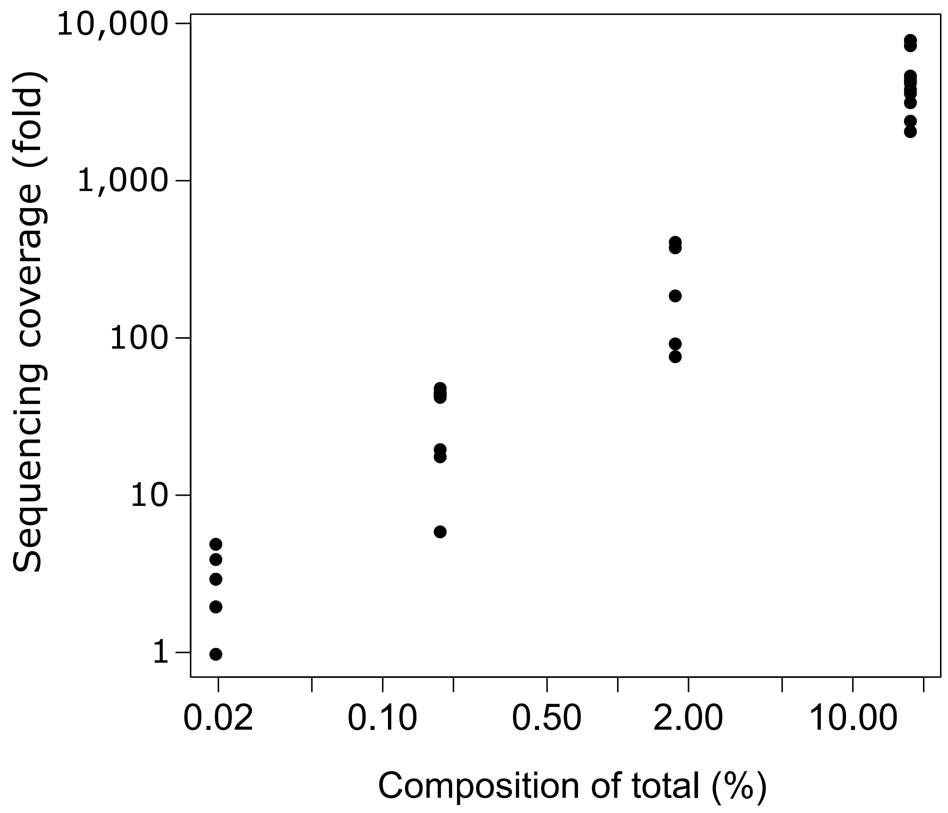

**Supplementary Figure 1.** Sequencing coverage vs. percent composition for the mock metagenome ATCC MSA-1003. Average sequencing coverage as mapped by minmap2 using the bacterial references listed in **Supplementary Table 1** is plotted against the expected relative composition of each bacterial species (percent of total).

### Supplementary Tables

#### Supplementary Table 1

| Sample | Strain | Composition (%) | Genome Reference* | Genome size (Mb) |
|---|---|---|---|---|
| *Acinetobacter baumannii* | ATCC 17978 | 0.18% | CP000521 | 3.97 |
| *Bacillus cereus* | ATCC 10987 | 1.80% | AE017194 | 5.22 |
| *Bacteroides vulgatus* | ATCC 8482 | 0.02% | CP000139 | 5.16 |
| *Bifidobacterium adolescentis* | ATCC 15703 | 0.02% | AP009256 | 2.09 |
| Clostridium beijerinckii | ATCC 35702 | 1.80% | NZ_CP006777 | 6.00 |
| *Cutibacterium acnes* | ATCC 11828 | 0.18% | CP003084 | 2.49 |
| *Deinococcus radiodurans* | ATCC BAA-816 | 0.02% | AE000513<br>AE001825<br>AE001827<br>AE001826 | 2.65<br>0.41<br>0.05<br>0.18 |
| *Enterococcus faecalis* | ATCC 47077 | 0.02% | NC_017316 | 2.74 |
| *Escherichia coli* | ATCC 700926 | 18..0% | U00096 | 4.64 |
| *Helicobacter pylori* | ATCC 700392 | 0.18% | AE000511 | 1.67 |
| *Lactobacillus gasseri* | ATCC 33323 | 0.18% | CP000413 | 1.89 |
| *Neisseria meningitidis* | ATCC BAA-335 | 0.18% | AE002098 | 2.27 |
| *Porphyromonas gingivalis* | ATCC 33277 | 18.0% | AP009380 | 2.35 |
| *Pseudomonas aeruginosa* | ATCC 9027 | 1.80% | PDLX01000000 | 6.34 |
| *Rhodobacter sphaeroides* | ATCC 17029 | 18.0% | CP000577<br>NC_009050 | 4.37 |
| *Schaalia odontolytica* | ATCC 17982 | 0.02% | DS264586<br>DS264585 | 2.39 |
| *Staphylococcus aureus* | ATCC BAA-1556 | 1.80% | CP000255<br>CP000256<br>CP000257<br>CP000258 | 2.87 |
| *Staphylococcus epidermidis* | ATCC 12228 | 18.0% | AE015929 | 2.50 |
| *Streptococcus agalactiae* | ATCC BAA-611 | 1.80% | AE009948 | 2.16 |
| *Streptococcus mutans* | ATCC 700610 | 18.0% | AE014133 | 2.03 |

*https://www.ncbi.nlm.nih.gov/nuccore/

**Supplementary Table 1.** Bacterial strain composition of metagenome staggered mix (ATCC® MSA-1003™).

### Supplementary Table 2

| Organism | Chromosome | Average Coverage (fold) |
|---|---|---|
| **Mus *musculus*** | 1 | 25 |
| **Mus *musculus*** | 2 | 25 |
| **Mus *musculus*** | 3 | 25 |
| **Mus *musculus*** | 4 | 25 |
| **Mus *musculus*** | 5 | 25 |
| **Mus *musculus*** | 6 | 25 |
| **Mus *musculus*** | 7 | 25 |
| **Mus *musculus*** | 8 | 25 |
| **Mus *musculus*** | 9 | 25 |
| **Mus *musculus*** | 10 | 25 |
| **Mus *musculus*** | 11 | 26 |
| **Mus *musculus*** | 12 | 25 |
| **Mus *musculus*** | 13 | 25 |
| **Mus *musculus*** | 14 | 25 |
| **Mus *musculus*** | 15 | 24 |
| **Mus *musculus*** | 16 | 24 |
| **Mus *musculus*** | 17 | 25 |
| **Mus *musculus*** | 18 | 24 |
| **Mus *musculus*** | 19 | 24 |
| **Mus *musculus*** | X | 23 |
| **Mus *musculus*** | Y | 0 |
| **Zea *mays*** | 1 | 22 |
| **Zea *mays*** | 2 | 22 |
| **Zea *mays*** | 3 | 22 |
| **Zea *mays*** | 4 | 22 |
| **Zea *mays*** | 5 | 22 |
| **Zea *mays*** | 6 | 22 |
| **Zea *mays*** | 7 | 22 |
| **Zea *mays*** | 8 | 22 |
| **Zea *mays*** | 9 | 22 |
| **Zea *mays*** | 10 | 22 |

**Supplementary Table 2**. Mapped coverage of the HiFi reads for the mouse and maize data sets reported across each chromosome.
